# The ZNF687 mutation of Paget’s disease associated with giant cell tumour causes severe bone remodelling alteration as a result of a deregulated osteoclast transcriptional program

**DOI:** 10.1101/2022.08.12.503488

**Authors:** Sharon Russo, Federica Scotto di Carlo, Giorgio Fortunato, Antonio Maurizi, Anna Teti, Danilo Licastro, Carmine Settembre, Fernando Gianfrancesco

## Abstract

Paget’s disease (PDB) is a late-onset bone remodelling disorder with a broad spectrum of symptoms and complications. One of the most aggressive forms is caused by the P937R mutation in the *ZNF687* gene. Although the genetic involvement of *ZNF687* in PDB has been extensively studied, the molecular mechanisms underlying this association remains unclear. Here, we describe the first *Zfp687* knock-in mouse model and demonstrate that the mutation recapitulates the PDB phenotype, showing a severe bone remodelling alteration. Through micro-computed tomography analysis, we observed that 8-month-old mice showed a mainly osteolytic phase, with a significant decrease in the trabecular bone volume affecting the femurs and the vertebrae of both heterozygous and homozygous mutant mice. In contrast, osteoblast activity was deregulated, beginning to produce disorganised bone. Noteworthy, this phenotype became pervasive in 16-month-old mice, where osteoblast function overtook bone resorption as the predominant event, as highlighted by the presence of woven bone in histological analyses, consistent with the PDB phenotype. Furthermore, we detected osteophytes and intervertebral disc degeneration, outlining for the first time the link between osteoarthritis and PDB in a PDB mouse model. Finally, we generated CRISPR-Cas9-based *Zfp687* knock-out RAW 264.7 cells, and noted a remarkable impairment of osteoclast differentiation capacity, reinforcing the relevance of Zfp687 during this process. RNA-sequencing on wild type and KO clones identified a set of genes involved in osteoclastogenesis under the control of Zfp687, i.e., *Tspan7, Cpe, Vegfc*, and *Ggt1*. Thus, this study established an essential role of Zfp687 in the regulation of bone remodelling, and may offer the potential to therapeutically treat PDB.

## Introduction

Paget’s disease of bone (PDB) is a late-onset skeletal disorder characterised by an impaired activity of bone remodelling, due to a high activity of bone degradation by osteoclasts followed by a disorganised bone deposition by osteoblasts^1,2^. PDB affects one (monostotic) or more (polyostotic) skeleton sites, and although any bone can be affected, there is a predilection for the skull, spine, pelvis, femur, and tibia. Bone pain, deformity, and pathological fractures, typically in the lytic phase of the disease, are the most common symptoms^1,2^.

A common complication of PDB is osteoarthritis (OA)^3^. Indeed, patients with PDB are more likely than age-matched controls to need total hip and knee arthroplasty, as a result of secondary OA^4^. A peculiar feature of OA is the formation of osteophytes, osteo-cartilaginous outgrowths that typically form at the joint margins, in the region where the synovium attaches to the edge of the articular cartilage and merges with the periosteum^5^. Osteophytes are established through growth of an initial cartilage template that is at least partially replaced with bone containing marrow cavities^6^. In PDB, the osteophyte occurrence is less clear. The original description of Sir James Paget in 1876 included the post-mortem analysis of a pathologic specimen of the femur of case 5, which demonstrated features of PDB at proximal and distal femur, with femoral head remodelling and osteophyte formation^7^. However, whether the formation of osteophytes is a prominent feature of PDB remains to be defined.

The most relevant PDB complication is the neoplastic degeneration of affected bones in osteosarcoma (OS/PDB) or giant cell tumour (GCT/PDB)^8^. Although rare, their occurrence, commonly seen in polyostotic PDB, is associated with more severe manifestations of the disease and reduced life expectancy. Osteosarcoma is the most fearful malignant transformation of PDB and shows an extremely poor prognosis that has not improved over the years, showing a 5-year survival rate almost nil, especially when metastasising to lungs^9^. In contrast, GCT rarely metastasises but is locally aggressive^10,11^. In the last decade, it has been proved that the distinct forms of the disease have different genetic basis^12–15^. The common form of PDB is due to mutations in the *SQSTM1* gene, encoding the p62 protein, involved in important cellular mechanisms, such as autophagy or the regulation of the NFκB signalling pathway^12,13^. Mutations in *SQSTM1* lead to an impaired and continuously activated NFκB pathway, resulting in more activated and multi-nucleated osteoclasts^16^. Two *Sqstm1* knock-in mouse models have been generated, harbouring the most common mutation (P392L), which only partially recapitulated the PDB phenotype: mutant mice developed osteolytic/osteosclerotic lesions predominantly affecting the long bones, while not involving the spine^17–19^. Interestingly, the most common anti-resorptive treatment with bisphosphonates prevented the formation of pagetic-like lesions in mutant animals^19^. PDB associated with giant cell tumour is instead negative for *SQSTM1* mutations and caused by a founder mutation (P937R) in the *ZNF87* gene. *ZNF687*-mutated patients present with a more severe clinical phenotype in terms of age of onset and number of affected sites^14^. Importantly, unlike *SQSTM1* mutation carriers, *ZNF687*-mutated patients show an inadequate response to bisphosphonates, and need massive doses of anti-resorptive drugs to cure the disease^20^. Moreover, it has been reported that pagetic patients harbouring the *ZNF687* mutation develop OA degeneration in 42.8% of cases, as PDB complication^10,14^. However, the role of *ZNF687* in bone metabolism needs to be further explored. Although *ZNF687* seems to be under the transcriptional regulation of NFκB, the pathway in which it is involved appears to be different than *SQSTM1*. To elucidate the molecular and pathological role of *ZNF687* in PDB, in this study we generated the knock-in mouse model carrying the P937R mutation in the homologous *Zfp687* gene, and performed its skeletal characterisation. This mouse model recapitulates the key clinical features observed in PDB patients, including polyostotic mixed osteolytic/osteosclerotic lesions but also an interesting link between PDB and OA, highlighted by the presence of osteophytes at the knee and vertebral joints in mice at 16 months of age (equivalent to about 55 human years). The role of Zfp687 in the pathogenesis of PDB was also confirmed in a CRISPR-based *Zfp687* knock-out RAW 264.7 cell model, displaying alterations of osteoclast differentiation process. Therefore, our study describes a novel mouse model that completely recapitulates the clinical manifestations of PDB, which not only enables to model the disease but also furthers our understanding of the role of Zfp687 in bone homeostasis.

## Results

### *ZNF687* is upregulated during human and murine osteoclastogenesis

Previously, we profiled the expression of *ZNF687* during the osteoclast differentiation of peripheral blood mononuclear cells (PBMCs) from healthy donors that pointed out a progressive upregulation of its expression^14^. Here, we first confirmed this upregulation at protein level during the physiological differentiation process in human and mice, confirming the key role of ZNF687 (**Fig. 1a, b**). Interestingly, *ZNF687*-mutated osteoclasts manifested a greater expression of *ZNF687* itself as compared to healthy individuals, supporting the gain-of-function status of the P937R mutation (**Fig. 1c, left**). Additionally, the expression of genes whose upregulation is crucial for successful osteoclastic differentiation (i.e., *TRAP, MMP-9, CTSK*) was also increased in mutation-bearing osteoclasts (**Fig. 1c, right**). This data agrees with the previous one demonstrating that patient-derived monocytes formed bigger and more active osteoclasts in response to pro-osteoclastic stimuli *in vitro*^20^. We therefore underline the relevance of ZNF687 in osteoclast biology, highlighting the positive modulatory effect on the osteoclastogenic process.

**Figure 1:**
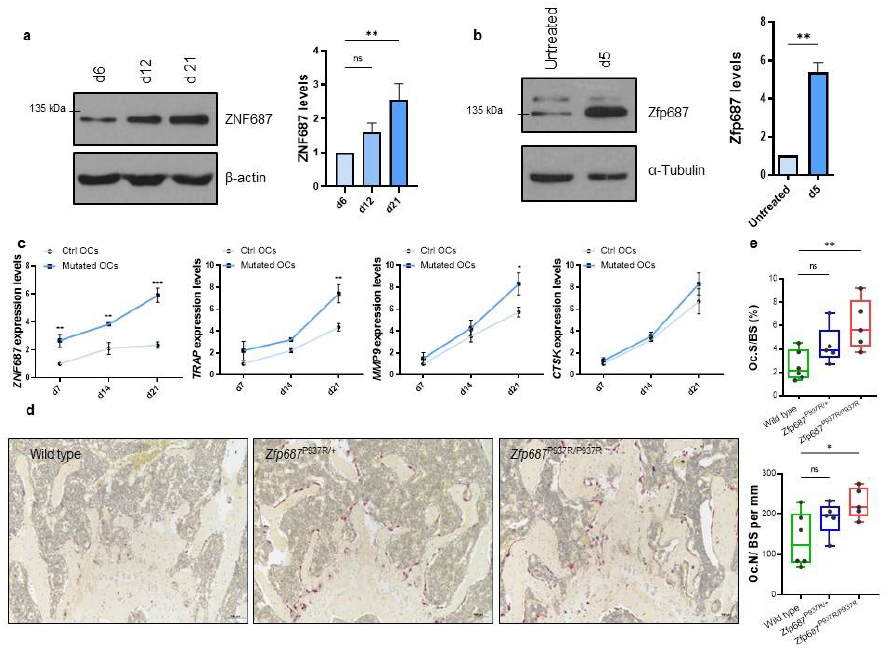
ZNF687 has a key role in osteoclastogenesis. **a)** Western blot detection (left) and densitometric quantification (right) of ZNF687 protein levels in differentiated osteoclasts derived from a healthy donor, upon sRANKL stimulation of PBMCs at day 6, 12, and 21. Western blots were normalised to b-actin. Data are presented as mean ± s.d. and statistical significance was assessed by one-way ANOVA with Dunnett’s multiple comparison test (**p < 0.01). **b)** Western blot detection (left) and densitometric quantification (right) of Zfp687 protein levels in differentiated osteoclasts derived from RAW 264.7 cells subjected to sRANKL stimulation for 5 days. Western blots were normalised to α-Tubulin. Data are presented as mean ± s.d. and statistical significance was assessed by unpaired t-test (**p < 0.01, two-tailed). **c)** Bar graphs showing gene expression analysis of *ZNF687* and osteoclastogenic markers *TRAP, MMP9*, and *CTSK* during osteoclast differentiation of WT and P937R-mutated PBMCs at day 7, 14, and 21. Data are presented as mean ± s.d. and statistical significance was assessed by two-way ANOVA with Sidak’s multiple comparison test (**p < 0.01; ***p < 0.001). **d)** Representative TRAP-stained sections of femoral growth plates of 3-month-old wild type, *Zfp687*^P937R/+^, and *Zfp687*^P937R/P937R^ mice showing osteoclastic activity in purple; nuclei are counterstained with haematoxylin. **e)** Box plots showing osteoclast surface to bone surface ratio (Oc.S/BS; top) and osteoclast number to bone surface ratio (Oc.N/BS; bottom) in *Zfp687*^P937R/+^ (n = 5) and *Zfp687*^P937R/P937R^ mice (n = 5) compared with wild type mice (n = 6) at 3 months of age. Data are presented as median ± s.d. and statistical significance was assessed by one-way ANOVA with Dunnett’s multiple comparison test (*p < 0.05).

### *ZNF687* mutation alters bone cell differentiation processes

To deepen the effect of the *ZNF687* mutation on bone metabolism, we generated a knock-in mouse model harbouring the c.2810C>G (P937R) mutation detected in GCT/PDB patients^14,20^. Both heterozygous and homozygous mice (referred to herein as *Zfp687*^P937R/+^ and *Zfp687*^P937R/P937R^, respectively) were viable and fertile, and the Mendelian distribution of genotypes in the litters was respected. Three-dimensional micro–computed tomography (µCT) evaluation of the trabecular bone composition of femurs and L4 lumbar vertebrae, as well as the cortical bone of femoral midshaft of 3-month-old mice were analysed, revealing neither skeletal abnormalities nor bone volume alterations in mutant animals (**Fig. S1, S2**). Even though at this stage there are no macroscopic differences, we observed a different bone cellular activity. First, femur sections were analysed for their tartrate-resistant acid phosphatase (TRAP) expression, revealing an increased osteoclast-dependent activity in mutant bones (**Fig. 1d**). Accordingly, we found significantly higher levels of both osteoclast surface (Oc.S/BS) and number (Oc.N/BS) per bone surface in histological sections of *Zfp687*^P937R/P937R^ mice, and a similar trend in *Zfp687*^P937R/+^ mice (**Fig. 1e**). Interestingly, we noted also an increased osteoblast number (Ob.N/BS) per bone surface within histological sections of mutant mice, especially homozygous mice (**Fig. 2a, b**). Taken together, these results underline the positive effect of the P937R mutation on number and activity of bone cells even before the overt phenotypic manifestation. If on one hand the effect of the *ZNF687* mutation on osteoclasts was previously explored^14,20^, on the other nothing is known about osteoblast differentiation. To fill this gap, we isolated bone marrow-derived mesenchymal stem cells (BM-MSCs) from 8-week-old wild type and homozygous mutant mice, which were then subjected to osteoblast differentiation. We first demonstrated a strong upregulation of *Zfp687* during the physiological differentiation process (**Fig. 2c**).

**Figure 2.**
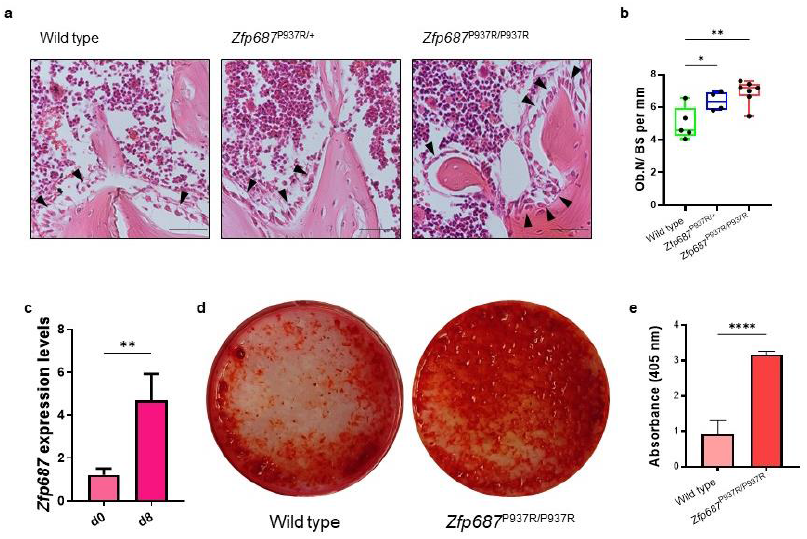
Zfp687^P937R^ differentiated osteoblasts display an increased mineralization potential. **a)** Representative H&E-stained images of proximal tibiae sections analysed for osteoblast (arrowheads) quantification in wild type, *Zfp687*^P937R/+^, and *Zfp687*^P937R/P937R^ mice at 3 months of age. Scale bars 50 µm. **b)** Box plots showing histomorphometry quantification of osteoblast number per bone surface ratio (Ob.N/BS) in wild type (n = 5), *Zfp687*^P937R/+^ (n = 4), and *Zfp687*^P937R/P937R^ (n = 7) mice at 3 months of age. Data are presented as median ± s.d. and statistical significance was assessed by one-way ANOVA with Dunnett’s multiple comparison test (*p < 0.05; **p < 0.01). **c)** Bar graph showing *Zfp687* expression analysis in wild type BM-MSCs either untreated (d0) or stimulated for 8 days towards osteoblastogenesis (d8). Data are presented as mean ± s.d. and statistical significance was assessed by unpaired t-test (**p < 0.01, two-tailed). **d)** Representative Alizarin Red Staining (ARS) images of wild type and *Zfp687*^P937R/P937R^ BMMSCs upon 8 days of osteogenic induction. **e)** Bar graph showing ARS quantification of wild type (n = 4) and *Zfp687*^P937R/P937R^ (n = 4) osteoblast differentiation upon 8 days of osteogenic induction. Data are presented as mean ± s.d. and statistical significance was assessed by unpaired t-test (****p < 0.0001, two-tailed).

Intriguingly, *Zfp687*^P937R/P937R^ osteoblasts manifested a remarkable higher capability of mineralisation and bone nodules formation in the presence of osteogenic factors (ascorbic acid and β-glycerophosphate), compared to wild type osteoblasts (**Fig. 2d**). Of note, mutant cells became fully differentiated osteoblasts as early as 8 days after osteogenic induction, while wild type cells displayed an expected slower differentiation process (**Fig. 2d, e**). We therefore conclude that the P937R mutation accelerates osteoblast formation and function, in agreement with what expected in bone remodelling alterations leading to PDB. During the histological analyses conducted on 3-month-old mice, we surprisingly noticed an increase in marrow adipocytes in mutant femurs. A mutual correlation between bone marrow adipose tissue (BMAT) and bone loss exists^21– 24^; yet, no research has been conducted to analyse this correlation in PDB. Therefore, we decided to quantify BMAT in the *Zfp687* mouse model. The distal tibiae of wild type and mutant mice at 3 months of age were stained with haematoxylin-eosin and subjected to measurement of the constitutive bone marrow adipose tissue (cBMAT). Our analysis highlighted that cBMAT was increased by ∼1.5 fold in both *Zfp687*^P937R/+^ and *Zfp687*^P937R/P937R^ mice (*p* = 0.0099 and *p* = 0.0079), as compared to wild type (**Fig. 3a, b**).

**Figure 3.**
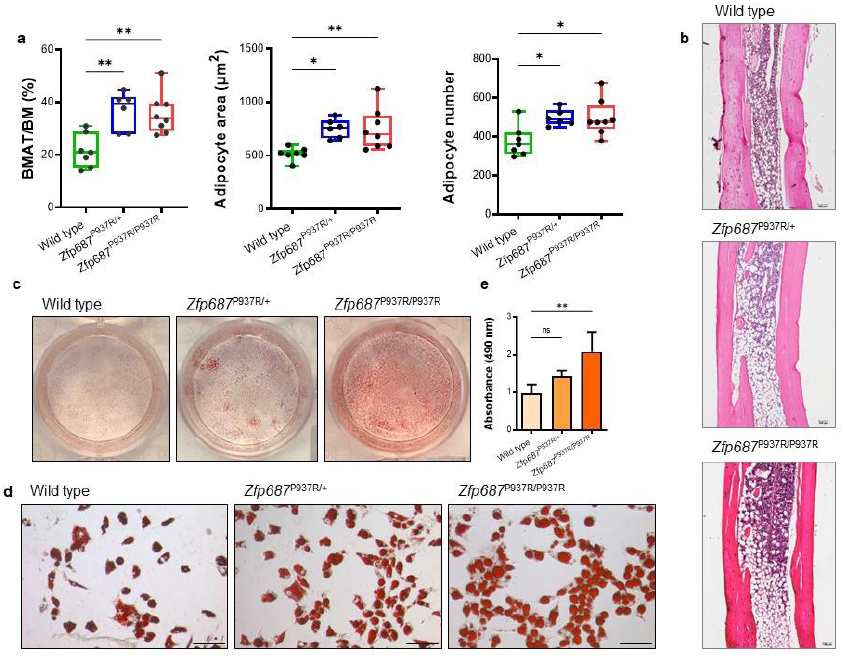
Zfp687 mutant mice display increased cBMAT composition at 3 months of age. **a)** Box plots showing the quantification of the tibial cBMAT volume expressed as the percentage of BMAT to total bone marrow ratio (BMAT/BM; left), the adipocyte area (middle), and the adipocyte number (right) of wild type (n = 7), *Zfp687*^P937R/+^ (n = 6), and *Zfp687*^P937R/P937R^ (n = 8) mice at 3 months of age. Data are presented as median ± s.d. and statistical significance was assessed by one-way ANOVA with Dunnett’s multiple comparison test (*p < 0.05; **p < 0.01). **b)** Representative H&E-stained images of distal tibia sections analysed for adipocytes measurements of the indicated genotypes. **c**, Oil Red O (ORO) staining of wild type, *Zfp687*^P937R/+^, and *Zfp687*^P937R/P937R^ BM-MSCs upon adipogenic induction and differentiation (plate view in **c)** and 20X magnification in **d))**. Scale bars 100 µm. Bar graphs show intensity of ORO staining (absorbance at 490 nm) of wild type (n = 4), *Zfp687*^P937R/+^ (n = 5), and *Zfp687*^P937R/P937R^ (n = 6) adipocytes. Data are presented as mean ± s.d. and statistical significance was assessed by one-way ANOVA with Dunnett’s multiple comparison test (**p < 0.01).

Both adipocyte number and area increased in mutant sections (**Fig. 3a**), indicating that mutant bone marrow contains more and larger fat cells. To confirm this observation, BM-MSCs were subjected to adipocyte differentiation *in vitro*, and as expected, mutant adipocytes appeared larger than wild type cells, and contained a huge amount of lipid droplets, as shown by ORO staining (**Fig. 3c-e**). Taken together, these data reveal that, in addition to the osteoclast differentiation program, the stromal cell commitment is also altered by the mutation in *Zfp687*.

### Adult Zfp687 mutant mice show trabecular bone loss and initial altered bone deposition

With the aim of shedding light on the early phase of Paget’s disease, we subjected to skeletal phenotyping mice at 8 months of age, which correspond to about 30 years in human, considering that the initial PDB diagnosis in *ZNF687*-mutated patients is generally around 45 years of age^14^. Parametric analysis revealed significant bone mass reduction affecting hind limbs and spine of mutant mice. In particular, the ratio of bone volume (BV) to total volume (TV; BV/TV) of femoral trabecular bone was decreased by 31% in *Zfp687*^P937R/+^ and 35% in *Zfp687*^P937R/P937R^ mice compared to wild type (*p* = 0.007 and *p* = 0.003, respectively) (**Fig. 4a, b**).

**Figure 4.**
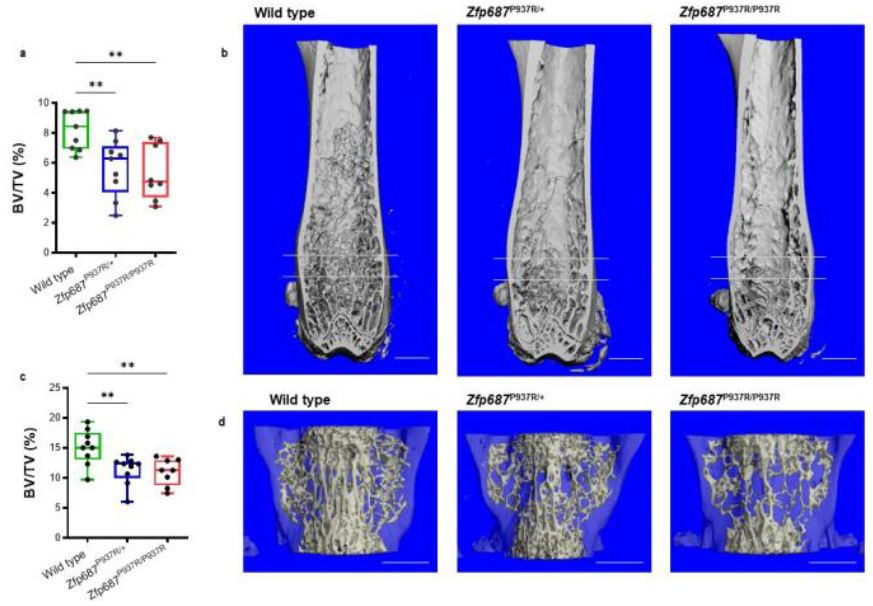
Zfp687 mutant mice show remarkable trabecular bone loss at appendicular and axial skeleton at 8 months of age. **a)** Box plots showing percentage of BV/TV by µCT in the trabecular bone of femoral distal epiphysis of 8-month-old wild type (n = 9), *Zfp687*^P937R/+^ (n = 9), and *Zfp687*^P937R/P937R^ (n = 8) mice. The region between the two gray lines represents the region of interest selected for the trabecular analysis. Data are presented as median ± s.d. and statistical significance was assessed by one-way ANOVA with Dunnett’s multiple comparison test (**p< 0.01). **b)** Representative µCT 3D reconstruction showing trabecular bone of femurs from wild type, *Zfp687*^P937R/+^, and *Zfp687*^P937R/P937R^ mice. Scale bars 1 mm. **c)** Box plots showing percentage of BV/TV by µCT in the trabecular bone of L4 vertebra of 8-monthold wild type (n = 9), *Zfp687*^P937R/+^ (n = 9), and *Zfp687*^P937R/P937R^ (n = 8) mice. Data are presented as median ± s.d. and statistical significance was assessed by one-way ANOVA with Dunnett’s multiple comparison test (**p< 0.01). **d)** Representative µCT 3D reconstruction of L4 vertebrae from wild type, *Zfp687*^P937R/+^, and *Zfp687*^P937R/P937R^ mice. Scale bars 1 mm.

The bone trabecular mass reduction was mainly due to a decrease in trabecular number (Tb.N) and, consequently, an increase in trabecular separation (Tb.Sp) (**Fig. S3a**). Additionally, trabecular bone mass in the spine was also reduced by 25% (*p* = 0.008) and 27% (*p* = 0.005) in the L4 vertebrae of *Zfp687*^P937R/+^ and *Zfp687*^P937R/P937R^ mice, respectively, with more sparse and thinner trabeculae than wild type littermates (**Fig. 4c, d, S3b**). Although neither significant cortical thickening (**Fig. 5a**), nor bone expansion, nor deformity were found at the midshaft of femurs of mutant animals, we sporadically observed the presence of lytic lesions affecting the cortical bone of long bones, in which TRAP-positive osteoclasts appeared giant-sized and multi-nucleated, as compared to the controls (**Fig. 5b**). A closer look at the osteoblast activity of 8-month-old mutant mice revealed that a misregulated bone deposition also occurred. In fact, histomorphometric analysis of femur sections through Von Kossa/Van Gieson staining showed an increased osteoid volume at both trabecular and cortical level in *Zfp687*^P937R/P937R^ mice that was compatible with an enhanced deposition of non-mineralised bone matrix (**Fig. 5c, d**). Collectively, these data indicate that the P937R mutation in 8-month-old mice leads to bone mass reduction and altered matrix deposition.

**Figure 5.**
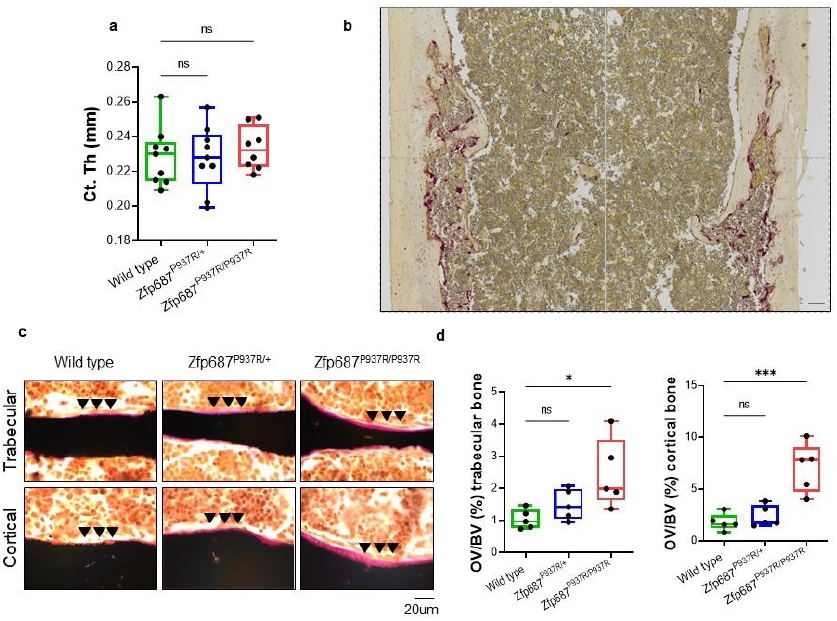
Severe impairment of bone remodelling in Zfp687 mutant mice at 8 months of age. **a)** Box plots showing quantitative measurements of cortical thickness of the femoral midshaft by µCT of 8-month-old wild type (n = 9), *Zfp687*^P937R/+^ (n = 9), and *Zfp687*^P937R/P937R^ (n = 8) mice. Data are presented as median ± s.d. and statistical analysis was assessed by one-way ANOVA with Dunnett’s multiple comparison test. **b)** TRAP staining of a femur section of a *Zfp687* mutant mice depicting a representative cortical osteolytic lesion. The figure shown is the result of a mosaic of 4 images of adjacent regions taken at 10x magnification. Scale bar 100 µm. **c)** Representative Von Kossa and van Gieson images of trabecular (upper) and cortical (lower) femur sections of wild type, *Zfp687*^P937R/+^, and *Zfp687*^P937R/P937R^ mice. Mineralised bone (black) and osteoid (pink) are visualised. Arrowheads indicate increased osteoid deposition. **d)** Box plots showing quantification of osteoid volume over bone volume percentage (OV/BV%) from Von Kossa-stained sections counterstained with van Gieson in trabecular (left) and cortical (right) bone of wild type (n = 5), *Zfp687*^P937R/+^ (n = 5), and *Zfp687*^P937R/P937R^ (n = 5) mice. Data are shown as median ± s.d. and statistical significance was assessed by one-way ANOVA with Dunnett’s multiple comparison test (*p < 0.05; **p < 0.01; ***p < 0.001).

### *Zfp687* mutation causes severe PDB in aged mice

Next, we performed skeletal phenotyping characterisation of 16-month-old mice, the equivalent of 55 human years, a state of full-blown pathology^14,20,25^. Remarkably, 87% of aged *Zfp687*^P937R/+^ and *Zfp687*^P937R/P937R^ mice developed polyostotic osteolytic-like lesions, affecting the lumbar spine and the calvarial bones, sites usually affected in pagetic patients (**Fig. 6a-c**). Three-dimensional reconstruction from µCT analyses also revealed the formation of large protruding osteophytes at the medial and lateral knee joint margins in 8 out of 16 mutant animals (**Fig. 6d**). These ectopic outgrowths, although less frequent were identified also at the spine (**Fig. 6e**) and made the movement of mice slow and difficult (**Supplementary Movie S1**). In fact, bi-dimensional µCT reconstruction highlighted enlargement of distal epiphysis of femurs and structural changes in the subchondral trabecular bone microstructure affected by the osteophyte formation (**Fig. 6f, arrowhead**). The occurrence of osteophytes together with vertebral fusion (shown in **Fig. 6a-b**) are compatible with an ongoing osteoarthritis process^26^.

**Figure 6.**
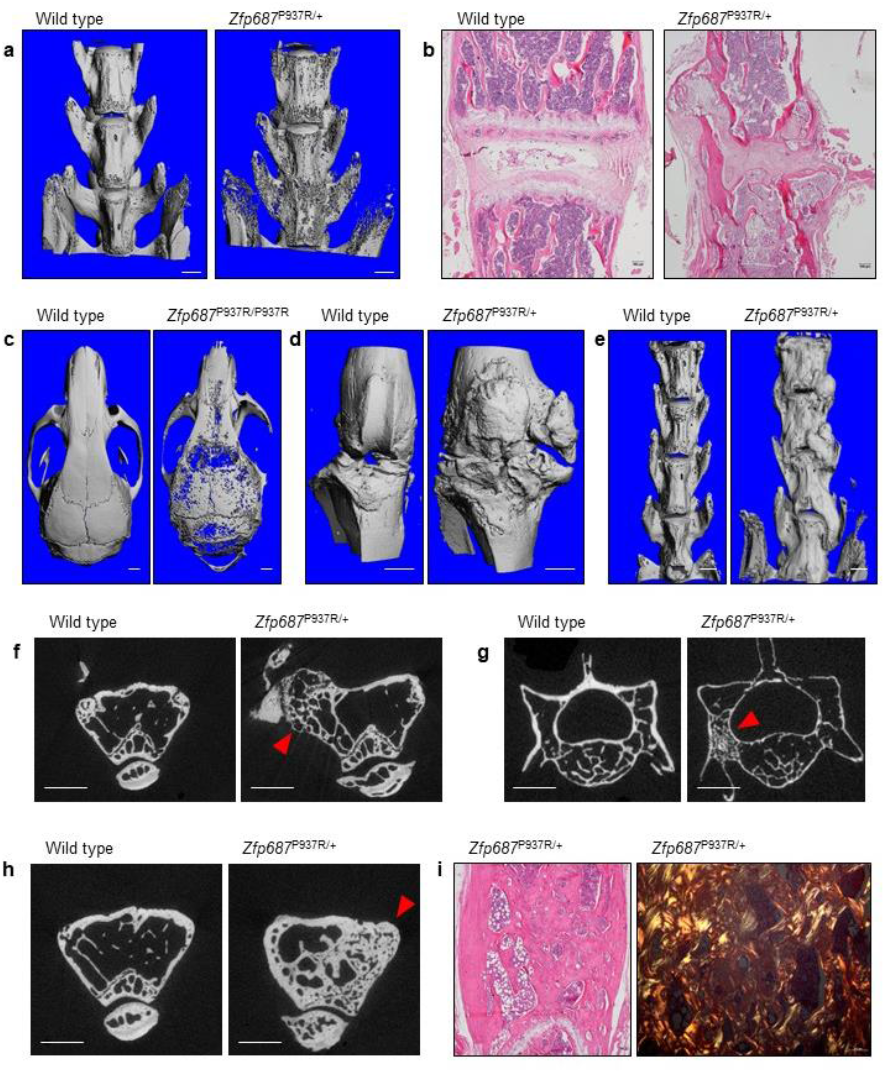
Aged Zfp687^P937R^ mutant mice develop pagetic lesions and osteophytes. **a)** Representative µCT reconstructed 3D images of the spine (lumbar vertebrae) of wild type and *Zfp687*^P937R/+^ mice, showing osteolytic cortical lesions and vertebral fusion in mutant mice at 16 months. **b)** Representative H&E-stained sections showing the intervertebral disk degeneration and cartilage degradation of the joint space between lumbar vertebrae of a *Zfp687*^P937R/+^ mutant animal compared to wild type (left). **c)** Representative µCT 3D images of osteolytic cortical lesions in calvarial bone of *Zfp687*^P937R/+^ mutant (right) compared to wild type (left). Scale bar 1 mm. **d, e)** Representative µCT 3D images of osteophyte formation at knee joint (in **d)**) and lumbar vertebrae (in **e)**) of wild type and *Zfp687*^P937R^ mutant mice. Scale bar 1 mm. **f)** Representative µCT cross-sections showing microarchitecture changes of subchondral bone (arrowhead) in correspondence of the osteophyte formation in the femur of *Zfp687*^P937R/+^ mutant compared to wild type. Scale bar 1 mm. **g)** Representative µCT cross-sections showing osteosclerotic lesions and ivory region (arrowhead) in L4 vertebra of a *Zfp687*^P937R/+^ mutant compared to wild type. Scale bar 1 mm. **h)** Representative µCT cross-sections showing chaotic structure and trabecularisation (arrowhead) of the femoral cortical bone in a *Zfp687*^P937R/+^ mutant, compared to wild type. Scale bar 1 mm. **i)** Histological sections of pagetic lesion in *Zfp687*^P937R/+^ femurs, showing the woven bone through H&E staining (left) and polarised-light microscopy (right).

Consistent with a pagetic phenotype, µCT scanning of 16-month-old mice also revealed osteosclerotic lesions in lumbar vertebrae (**Fig. 6g**) and distal epiphyses of femurs (**Fig. 6h**) of mutant mice. We detected enlarged bones with ivory regions, and trabecularisation of cortical bone (**Fig. 6h, arrowhead**). Histological analysis of these lesions showed increase in bone resorption and formation with accumulation of woven bone, as detected by polarised light microscopy (**Fig. 6i**). The frequency and type of skeletal defects detected in aged mice is reported in **Table 1**. Thus, altogether these results illustrate that the P937R mutation is necessary and sufficient to fully develop a severe form of PDB.

**Table 1.** Phenotypic analysis of skeletal alterations in 16-month-old mice.

|  | osteolysis in<br>appendicular<br>skeleton | osteolysis in<br>axial<br>skeleton | osteosclerosis in<br>appendicular skeleton | osteosclerosis in<br>axial skeleton | osteophyte<br>at knee joint | osteophyte<br>at spine | at least 2 sites<br>affected | at least 3 sites<br>affected |
| --- | --- | --- | --- | --- | --- | --- | --- | --- |
| Wild type | 0% | 11% | 0% | 0% | 0% | 0% | 0% | 0% |
| *Zfp687*<sup>P937R/+</sup> | 22% | 70% | 11% | 20% | 67% | 30% | 80% | 45% |
| *Zfp687*<sup>P937R/P937R</sup> | 40% | 60% | 0% | 0% | 0% | 0% | 60% | 20% |

### Zfp687 is an essential driver of osteoclast differentiation

To obtain mechanistic insights on the role of the *Zfp687* gene in bone metabolism, we induced the CRISPR/Cas9-mediated *Zfp687* knock-out in the murine RAW 264.7 macrophage cell line. We selected three different heterozygous clones (*Zfp687*^*+/-*^). TRAP staining performed after 5 days of sRANKL stimulation revealed that osteoclast formation and differentiation were severely impaired in *Zfp687*^*+/-*^ cells (**Fig. 7a**). Indeed, the number of mature osteoclasts, identified as TRAP-positive cells with more than 3 nuclei, dramatically decreased by an average of 73% in all KO clones analysed, compared to the wild type counterpart (**Fig. 7b**). Moreover, we observed that *Zfp687*^*+/-*^ osteoclasts showed strongly reduced surface area (average 3800 µm^2^), while wild type cells were typically larger (average 11000 µm^2^) (**Fig. 7c**). This result indicates that a single copy of the Zfp687 transcription factor is not sufficient to drive a proper osteoclastogenesis upon sRANKL stimulation.

**Figure 7.**
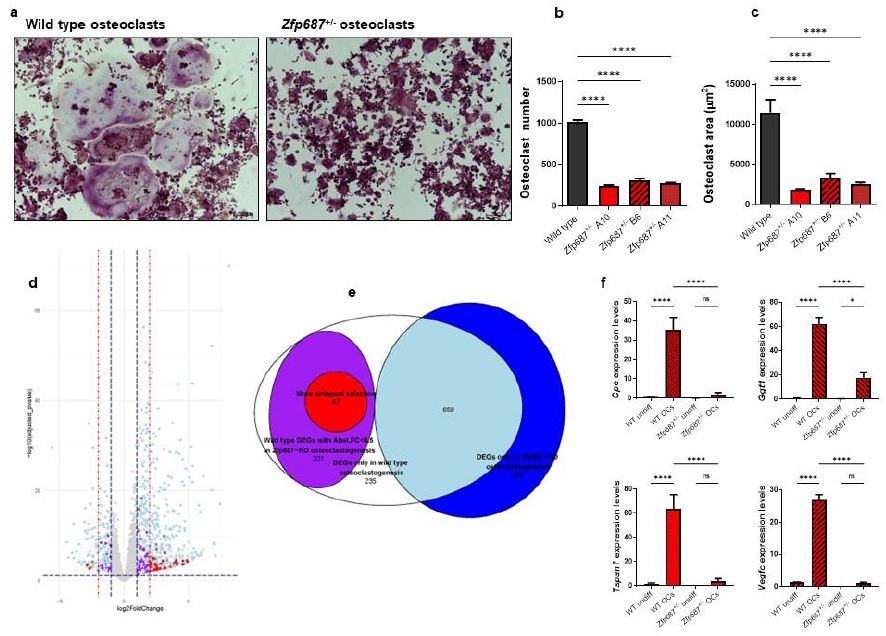
The pivotal role of ZNF687 for a proper osteoclast differentiation. **a)** Representative images of TRAP-stained osteoclasts from wild type and *Zfp687*^+/-^ RAW 264.7 cells after 5 days of sRANKL stimulation. **b)** Bar graph showing the mean number of TRAP+ osteoclasts (more than 3 nuclei/cell) in wild type (n = 2) and *Zfp687*^+/-^ (n = 6) clones. Data are presented as mean ± s.d. and statistical significance was assessed by one-way ANOVA with Dunnett’s multiple comparison test (****p < 0.0001). **c)** Bar graph showing the mean area of TRAP^+^ osteoclasts in wild type and *Zfp687*^+/-^ clones. **d)** Volcano plot showing the distribution of differentially expressed genes (DEGs) between wild type RAW 264.7 and *Zfp687*^*+/-*^ clones. In light blue, DEGs shared by wild type and *Zfp687*-KO osteoclastogenesis; in purple, DEGs in wild type but unchanged in *Zfp687*-KO osteoclastogenesis (log2foldchange ≥ 1 and ≤ - 1); in red, selected DEGs in wild type but unchanged in *Zfp687*-KO osteoclastogenesis (log2foldchange ≥ 2 and ≤ −2). **e)** The Venn map of DEGs in wild type and *Zfp687*-KO osteoclastogenesis. **f)** Bar graphs showing the relative expression of genes selected for RNA-seq validation. Undiff: undifferentiated; OCs: osteoclasts. Data are presented as mean ± s.d. and statistical significance was assessed by one-way ANOVA with multiple comparison test (*p < 0.05; ****p < 0.0001).

To identify Zfp687 target genes that may influence the correct process of osteoclast differentiation, we performed RNA-sequencing on RNA extracted from wild type RAW 264.7 and three *Zfp687*^*+/-*^ clones, before and after RANKL osteoclastogenic induction for 5 days. Under physiological condition, i.e. in wild type extracts, a total of 1322 genes were differentially expressed during the process of osteoclast differentiation: 1055 upregulated and 267 downregulated genes (**Fig. 7d**). The CLEAR (Coordinated Lysosomal Expression and Regulation) Signalling, which regulates lysosomal biogenesis and function, was the most upregulated pathway (*p* = 5.24E-10)^27^. Conversely, transcriptomic profiling of osteoclast differentiation process in *Zfp687*^*+/-*^ cells highlighted that 381 genes, previously detected as upregulated in the wild type context, remained instead unchanged (**Fig. 7e**). Similarly, 47 genes previously found as downregulated during the control osteoclastogenesis, were unchanged in the three mutant processes (**Fig. 7d**). This data suggests that these genes, whose expression was unperturbed by RANKL stimulation in *Zfp687*^*+/-*^ cells, might be under the transcriptional control of Zfp687. We observed that the crucial genes for osteoclastogenesis, including *Cathepsin K* (*CTSK*), *Tartrate-resistant acid phosphatase* (*TRAP*), and *Matrix metalloproteinases-9* (*MMP-9*), were significantly upregulated in both control and mutated processes confirming the phenotypic evidence that, mutant osteoclastogenesis was severely impaired but not completely abolished. Therefore, we focused our attention on the genes that remained unchanged in *Zfp687*^*+/-*^ cells after stimulation. By using a more stringent cut-off parameter we restricted our analysis to 92 upregulated and 5 downregulated genes (**Fig. 7e)**. From this list, we selected the most differentially expressed genes and those with a clear involvement in osteoclast differentiation, obtaining a high-ranking list of 16 genes (15 up- and 1 downregulated) (**Table 2**). Among them, *Tspan7, Cpe, Vegfc*, and *Ggt1* were previously characterised as having a role in osteoclastic differentiation^28–34^. We confirmed through real time PCR that their expression only increased in physiological osteoclastogenesis, and remained unvaried in stimulated *Zfp687*^*+/-*^ cells (**Fig. 7f**). In conclusion, we demonstrated that the Zfp687 transcription factor is a crucial regulator of osteoclast differentiation by regulating several important genes whose study could allow to discover further molecular mechanisms at the base of the PDB.

**Table 2.**
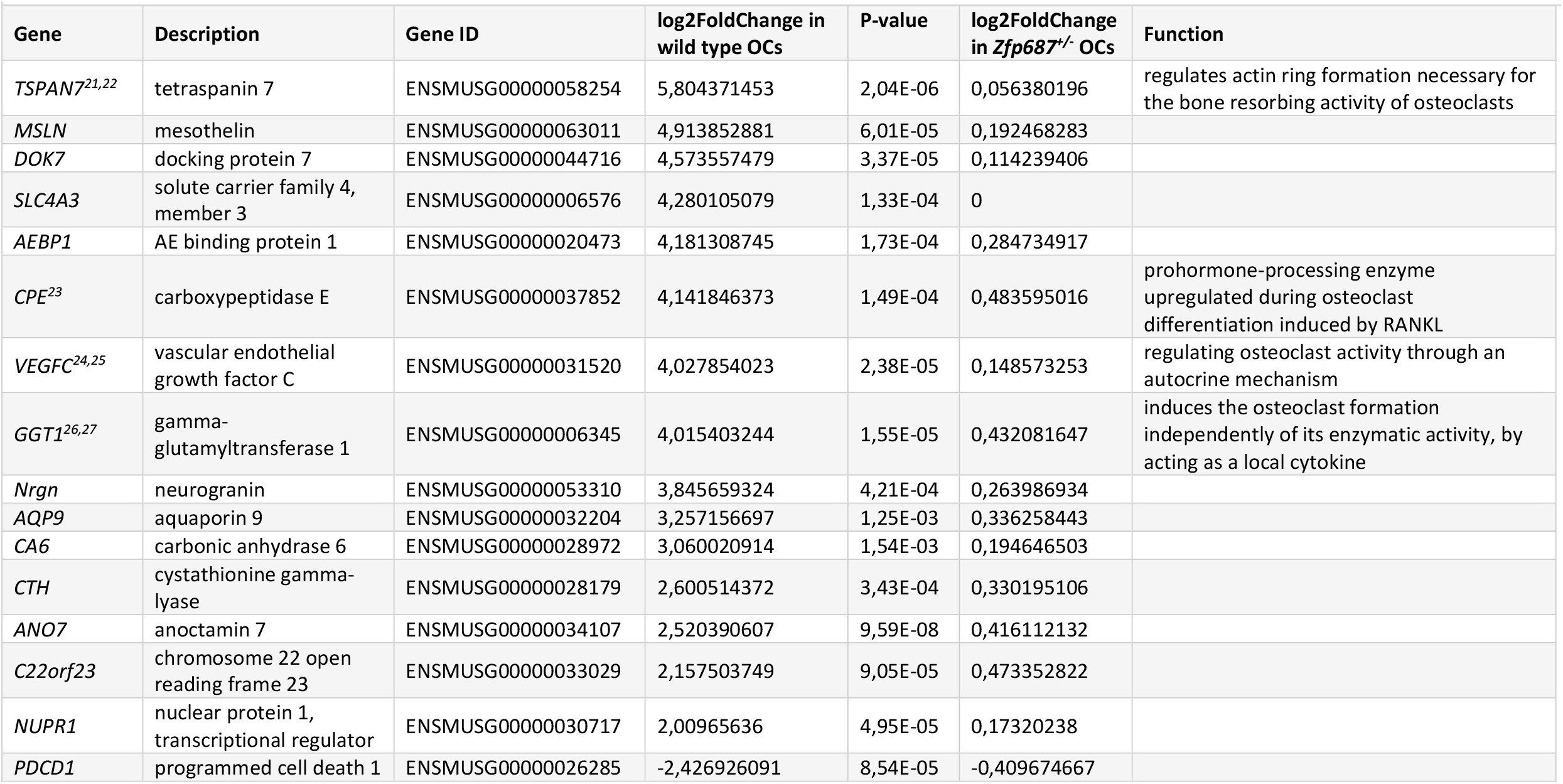
Information of 16 differential expressed osteoclastic genes associated with the loss of *Zfp687* gene.

## Discussion

Paget’s disease of bone (PDB) is a bone remodelling disorder, in which osteoclasts appear giant-sized with an increased bone destruction activity, and osteoblasts follow this process through their disorganised bone deposition activity^1^. As a result, the remodelled bone becomes weaker, deformed and more likely to fracture. Previously, we described that the founder P937R mutation in the *ZNF687* gene is causative of a severe form of PDB complicated by giant cell tumour degeneration (GCT/PDB)^14,20,35^. In order to determine the effect of the P937R mutation on bone metabolism and PDB pathogenesis, we generated the *Zfp687* knock-in mouse model. Mice harbouring the mutation, in heterozygosity and homozygosity, develop impressive skeletal phenotype that worsen as they age, mirroring the late onset of PDB in humans. Specifically, µCT analyses performed on adult and aged mutant mice displayed bone remodelling alteration starting from 8 months of age, affecting both the axial and the appendicular skeleton. This phenotype was highly pervasive at 16 months of age, in agreement with the age of appearance of full-blown disease in human patients.

Intriguingly, no P937R homozygous patient has ever been found in our cohort of PDB individuals, leading to the hypothesis that the homozygosity for the mutation could be lethal. Nonetheless, homozygous mutant mice are viable and do not show a more aggressive phenotype than heterozygous animals, indicating that the human mutation is so rare that the chance for an individual of inheriting two mutated alleles is highly unlikely. This observation also points out that the inheritance of a single mutated allele is necessary and sufficient to drive the disease.

As PDB is a genetically heterogeneous diseases, other two PDB mouse models have been described so far, harbouring the most common *Sqstm1* mutation (P394L) found in human patients^17–19^. However, these mutant mice only partially recapitulate the PDB phenotype, developing few bone alterations exclusively at the appendicular skeleton, not earlier than 18 months of age. This is presumably due to the pleiotropic role of p62, the multifunctional protein encoded by *Sqstm1*, which is involved in multiple cellular functions, such as clearance of mis-folded protein, autophagy, cell survival, regulation of Keap1–Nrf2 and NFκB pathways^36^. On the other side, the exact function of ZNF687 is not completely understood. The *Zfp687* mouse model, however, developed a much more severe bone phenotype, fully replicating the occurrence of osteolytic, mixed osteolytic/osteosclerotic, and osteosclerotic phases of Paget’s disease. Furthermore, unlike the *Sqstm1* mutant mice, here we report that *Zfp687* mutants display bone remodelling alterations at the spine, enabling researchers to better model and study the disease. In these mice, the disease is so severe that additional complications occur. We indeed found osteophytes at the knee joints and vertebral fusion in 50% of aged mutant mice as a consequence of degeneration in osteoarthritis (OA). In fact, OA is a quite common complication of PDB, which becomes even more frequent in *ZNF687*-mutated patients (>40% of cases)^3,14^. In our mouse model, we also highlighted an unexpected increase in BMAT in both heterozygous and homozygous knock-in mice. Two different studies reported independently that BMAT-derived RANKL induces osteoclastogenesis and bone remodelling, indicating that excessive RANKL generated by bone marrow adipocytes is an underlying cause of skeletal disorders^37,38^. Therefore, we cannot exclude a positive effect of the mutation on BMAT that further boosts osteoclast differentiation and therefore results in a more aggressive phenotype. Our data also indicated that BM-MSCs harbouring the P937R mutation are capable to differentiate towards both osteoblast and adipocyte cell lineage with a higher efficiency than control cells. To explain this apparently counterintuitive phenomenon, which counteracts the mutually exclusive differentiation program described for fate-decision in one of the two different lineages^39^, we could speculate that bone marrow of mutated mice might be enriched either in progenitors with both osteo- and adipo-differentiation properties or in MSCs with an increased proliferation ability. Thus, additional studies are needed to better understand the involvement of Zfp687 during the BM-MSCs fate commitment and differentiation.

In this study, we also reported a set of genes involved in osteoclastogenesis under the control of Zfp687. Among them, *Tspan7, Cpe, Vegfc*, and *Ggt1* are described as having a role in regulating the bone-resorbing function of osteoclasts^28–34^. Of course, additional functional studies are necessary to disclose the specific effect of these genes on PDB pathogenesis or to identify additional molecular mechanisms. Finally, an open question related to this model remains the development of the tumour. PDB patients with the *ZNF687* mutation, if untreated, undergo giant cell tumour degeneration. Up to 24 months of age, this model did not develop any bone tumours. Nonetheless, the evidence that mutant of Zfp687 did not drive tumorigenesis within 24 months of age is not surprising, mainly in animals with functional p53. It may be interesting to verify the tumorigenic capacity of Zfp687 mutation in a p53-deficient background. However, mutant mice still seem to be predisposed to tumorigenesis, because 2 out of 6 heterozygous and 5 out of 11 homozygous mutant mice at 20 months of age developed hepatic nodules resembling tumour masses (data not shown). This result was consistent with the role for ZNF687 as an oncogene found in hepatocellular carcinomas^40^

## Acknowledgements

The research leading to these results has received funding from AIRC under IG 2020 - ID. 25110 project – P.I. F.G.

## Author contributions

S.R., F.S.d.C., and F.G. conceived the study and wrote the paper, drafted the figures and tables. S.R. and F.S.d.C. carried out in vitro and ex vivo experiments, and performed statistical analyses. F.S.d.C., S.R., and F.G. analysed the results. D.L. performed analyses of RNA sequencing data. A.M. and A.T. performed Von Kossa staining and data analyses. G.F. performed initial analysis on RAW 264.7. C.S. helped in µCT study and data analysis. F.G. supervised the study and data analysis. All authors read and approved the final version of the manuscript.

## Conflict of interests

The authors declare no competing interests.

## Data availability

All relevant data supporting the key findings of this study are available within the article. Raw sequencing data are available upon request.

## Materials and Methods

### Generation of the Zfp687^P937R^ knock-in mouse model

To generate the PDB mouse model carrying the P937R mutation in the *Zfp687* gene, we adopted the homologous recombination strategy. A BAC library was used as a template to amplify the *Zfp687* locus for the homologous recombination. PCR fragments were cloned into the pGND vector as long and short homology arms. The P937R mutation (c.C2810G) was introduced in the long homology arm by PCR-mediated mutagenesis (Quik-Change Lightning, Agilent), using the following primers: sense 5’-GTT-GGTCGGGGTCGCTCAGGGGAGC-3’; anti-sense 5’-GCTCCCCTGAGCGACCCCGACCAAC-3’. Targeting of the construct was done into the E14Tg2a embryonal stem (ES) cell line, and targeted ES cell clones were identified by Southern blot analysis using a 3’ probe and an internal Neo probe on SphI-cut genomic DNA, and a 5’ probe on BglII-cut genomic DNA. Sanger sequencing confirmed the presence of the c.C2810G mutation. One *Zfp687*-mutated ES clone was injected into C57BL6 blastocysts to establish a mutant mouse colony. The neomycin resistance gene was removed crossing the mice with a Cre transgenic mouse. These mice were maintained on a mixed genetic background. Genotyping was performed with allele-specific primers: forward 5’-GACAGCCCTCTAAACCTCAAGACC-3’; reverse 5’-AGCAGGAGCATTAGTGTTGGATTC-3’, leading to the amplification of two different sized products depending on the presence or absence of the loxP site. Animals were handled in accordance with the authorization no. 125-2021-PR released by the Italian Ministry of Health; all mice were housed in a pathogen-free barrier environment.

### Micro-Computed Tomography (μCT) analysis

Male mice were sacrificed by CO_2_ inhalation at the indicated age, and the skin was removed. Femurs, tibiae, spine, and skull were cleaned from adherent and soft tissue, fixed in 4% paraformaldehyde, PFA (Sigma-Aldrich, #158127) for 24 h at 4 °C, and then stored in 70% ethanol. μCT analyses were conducted using the SCANCO Medical-μCT40 (Scanco Medical AG, Bassersdorf, Switzerland). For bone morphometry, femoral trabecular and cortical bone were scanned using the following parameters: E = 70 kV; I = 114 μA; integration time of 600 ms; 3 μm isotropic voxel size. Femurs were scanned 1 mm from the distal boarder of the growth plate. Cortical thickness was measured at the midshaft region of the femur diaphysis. For femur trabecular bone and for midshaft cortical bone, 209 and 36 slices were respectively analysed. Vertebral trabecular bone was analysed by selecting and scanning the whole L4 vertebra, using the last rib-bearing thoracic vertebrae as reference, using the following parameters: E = 55 kV; I = 145 μA; integration time of 300 ms; and 12 μm isotropic voxel size. The trabecular bone of the vertebral body was evaluated immediately below the superior growth plate. A lower threshold of 270 was used for the evaluation of all scans. For trabecular bone of femur and L4 vertebra, the structural parameters bone volume/total volume (BV/TV), trabecular thickness (Tb.Th), trabecular number (Tb.N), and trabecular separation (Tb.Sp) were considered. To reconstruct solid 3D images, selected bone samples were scanned in high resolution. Femur, lumbar spine, and skull samples were scanned using the following parameters: E = 70 kV; I = 114 μA; integration time of 300 ms; 6 μm isotropic voxel size. The reconstructed solid 3D images were applied for visualizing bone morphology and microarchitecture.

### Histological analysis

For histology, long bones (femurs and tibiae) and the lumbar region of the spine were decalcified in 14% ethylenediaminetetraacetic acid, EDTA (Sigma-Aldrich, #27285) for 14 days, replacing the solution every 3 days. Then, bone samples were dehydrated with ethanol series (70, 80, 90, 100% Et-OH), treated with xylene, and paraffin-embedded. Bone slices of 3 and 5 μm were obtained by manual microtome. Bone sections were subjected to wax-removal procedure by xylene treatment and rehydration through a graded series of alcohol (100, 90, 80, 70% Et-OH) and tap water. Bone sections were stained with haematoxylin and eosin (H&E) for general tissue morphology, and with tartrate-resistant acid phosphatase (TRAP) (Sigma-Aldrich, #387A) to detect osteoclasts activity, according to standard protocols. After staining, bone sections were dehydrated, mounted with a mounting medium (Bio-Optica, #05-BMHM), covered with a coverslip and analysed by transmission light microscopy, using Nikon Motorized Eclipse Ni-U Microscope and Nikon Manual Optical Microscope. Osteoclasts surface and number were measured by the TRAPHisto open-source software^41^. For osteoblast quantification (Ob.N/BS), paraffin-embedded tibiae samples were stained with H&E. Osteoblasts were identified as cuboidal cells lining the trabecular bone surface in the proximal metaphyseal region of tibiae. Osteoblasts were manually counted from 5 fields, 2 slices per animal (20X magnification), and the mean was calculated for each animal (n

= 5 wild type; n = 4 *Zfp687*^P937R/+^; n = 7 *Zfp687*^P937R/P937R^). Bone surface was determined by ImageJ software. For BMAT quantification and adipocyte measurements, H&E staining was performed on tibiae. Adipocyte cells were identified by their thin cytoplasmic layer that lines the lipid droplet and forms the ghost-like remnant of the adipocytes. Adipocyte ghost cells were manually counted from at least 3 sections spaced 100 µm per animal (4X magnification) (n = 7 wild type; n = 6 *Zfp687*^P937R/+^; n = 8 *Zfp687*^P937R/P937R^). All the adipocytes in each section were counted and 160 adipocytes/section were measured for the area evaluation. All the measurements were determined using ImageJ software.

### Von Kossa/Van Gieson staining and osteoid quantification

For osteoid and matrix mineralisation evaluation, 7 µm sections from MMA-embedded femurs of 8-month-old mice were stained with Von Kossa and counterstained with Van Gieson, according to standard procedures. Briefly, aqueous silver nitrate solution was added on the slides, which were then incubated with soda-formol solution for 5 minutes. Then, sodium thiosulfate was added and let incubate for 5 minutes to remove unreacted silver. Von Kossa stained samples were rinsed in tap water and counterstained with Van Gieson solution for 30 minutes. Slices were mounted and covered with a coverslip and analysed by transmission light microscopy. Osteoid quantification was performed using the threshold colour function of ImageJ software.

### Generation of Zfp687 knock-out RAW 264.7 cell clones using CRISPR-Cas9 technology

*Zfp687* knock-out RAW 264.7 cell clones were obtained by the CRISPR-Cas9 technology. The small guide RNA (sgRNA) was designed by using the tool at http://crispor.tefor.net/, targeting the exon 2 at the fourth ATG, predicted as surrounded by a Kozak consensus sequence (sense 5’-CAC-CGCCTCAAGGGGCCTTGAAAC-3’). The sgRNA was cloned in the pSpCas9(BB)-2A-GFP plasmid (Addgene, #48138). Then, the genetic transformation of RAW 264.7 cells was obtained by nucleofection using the Amaxa Cell Line Nucleofector Kit V (Lonza) for RAW 264.7 and following the protocol for Amaxa Nucleofector. After 24 h, single GFP-positive cells were sorted in 96-well plates with Becton Dickinson FACS AriaIII. We obtained three heterozygous *Zfp687* knock-out clones; homozygous knock-out clones were never detected. The clones harbouring distinct heterozygous frameshift mutation (*Zfp687*^*+/-*^) were confirmed by Sanger sequencing.

### Cell culture

Primary murine bone marrow-derived mesenchymal stromal cells (BM-MSCs) were obtained from femurs and tibiae of 8-weeks-old mice, adapted from the method described in^42^. Briefly, mice were euthanized through CO_2_ and immediately after the sacrifice, femurs and tibiae were carefully cleaned of all the connective tissues; both the distal and proximal ends of bones were cut, and the marrow was centrifuged out. ACK (Ammonium-Chloride-Potassium) lysing buffer was used to eliminate red blood cells. Total bone marrow cells were cultured in complete expansion medium (MesenCult Expansion Kit (Mouse) #05513, Stem Cell Technologies), at 37 °C, 5% CO_2_. BM-MSCs were expanded for 7 days. For the osteogenic differentiation, cells were detached using 0.25% Trypsin-EDTA, plated in 24-well plates and cultured in complete expansion medium until they reached 80-90% confluency. Then, to induce osteogenic differentiation, medium was replaced with complete MesenCult Osteogenic Medium (MesenCult Osteogenic Stimulatory Kit (Mouse) #05504, Stem Cell Technologies), and cells were cultured at 37 °C, 5% CO_2_. Medium was changed every 3 days for 8 days. Differentiated osteoblasts were fixed in 70% ethanol and stained with Alizarin Red Solution (Sigma-Aldrich #A5533). De-staining was conducted to quantitatively determine mineralisation by adding acetic acid. Absorbance was measured in the microplate reader PerkinElmer luminometer (Victor X3) at 405 nm.

For the adipogenic differentiation, medium was replaced with MesenCult Adipogenic Differentiation Medium (Mouse) #05507 (Stem Cell Technologies) for 6 days. Adipogenic induction was assessed by Oil Red O (ORO) staining (Sigma-Aldrich #O1392), following the manufacturer’s instruction. For quantification, ORO was extracted by adding isopropanol, and absorbance was read in the microplate reader PerkinElmer luminometer (Victor X3) at 490 nm.

RAW 264.7 cells were cultured in DMEM High Glucose GlutaMAX (Gibco), with 10% FBS, 1% penicillin/streptomycin, and 1% L-Glutamine at 37 °C, 5% CO_2_. For osteoclast differentiation, 5×10^3^ cells were plated in 24-well plates and the medium was switched in Minum Essential Medium α (MEM-α) GlutaMAX (Gibco), with 10% FBS, 1% penicillin/streptomycin, and 1% L-Glutamine. The day after, medium was changed and supplemented with 100 ng/ml sRANKL (Peprotech) for the osteoclastogenic induction. The medium was changed every 48 h, until the end of the differentiation (5 days upon stimulation). Differentiated osteoclasts were fixed in 4% PFA and stained with tartrate-resistant acid phosphatase (TRAP) (Sigma-Aldrich).

### Protein extraction and Western blotting

Total protein extraction from murine cell lines (RAW 264.7 cells and osteoclast-derived cells) was performed in RIPA buffer (50 mM Tris–HCl pH 7.5; 150 mM NaCl; 1 mM DTT; 50 mM sodium fluoride; 0.5% sodium deoxycholate; 0.1% SDS; 1% NP-40; 0.1 mM phenylmethanesulfonylfluoride; 0.1 mM sodium vanadate) with 1X proteinase inhibitor cocktail (Applied Biological Materials #G135). Protein quantification was obtained by the Bradford method (Bio-Rad #5000006). Protein samples were boiled at 95 °C for 5’ and then were separated by SDS-PAGE electrophoresis, using 8-16% Tris-Glycine gels (Invitrogen #XP08160BOX). Samples were transferred on a nitrocellulose membrane (Invitrogen #IB23002), blocked with 4% w/v non-fat dry milk dissolved in TBS-T (1X TBS, 0.05% Tween-20) for 1 h at RT. Primary antibodies used for the Western blot experiments were rabbit anti-ZNF687 (1:3’000, Novus NBP2-41175), mouse anti-β-Actin (1:10’000, SantaCruz #47778), and mouse anti-α-Tubulin (1:15’000, Sigma-Aldrich #T6074). Membranes were incubated with secondary antibodies conjugated with HRP for 1 h at RT. The bands were visualised using enhanced chemiluminescence detection reagents (Advansta #K-12043-D10) and autoradiographic films (Aurogene #AU1101). Equal loading was confirmed by using antibody against anti-β-Actin and α-Tubulin. The intensity of the western blot signals was determined by densitometry analysis using the ImageJ software and normalised to the density value of the loading control. Protein extracts of peripheral blood mononuclear cells (PBMCs) and differentiated osteoclasts from a healthy donor and P937R-mutated patient were previously collected^20^ and already present in our laboratory.

### RNA isolation and qRT-PCR analysis

Total RNA extraction (from BM-MSCs, mouse osteoblasts, RAW 264.7 cells, and differentiated osteoclasts) was obtained using TRI-Reagent (Sigma-Aldrich #T9424), following the manufacturer’s instruction. One microgram of total RNA was retrotranscribed in cDNA using the RevertAID RT kit (ThermoFisher #K1622). qRT-PCR was performed using the SYBR Select Master Mix for CFX (Applied Biosystems) and specific primers listed in **Table 3** on CFX Opus RT PCR System instrument. The transcript levels were normalized to the levels of *Gapdh* within each sample, and the DDCT method was used. The reaction was conducted in triplicate. cDNA of PBMCs and differentiated osteoclasts derived from healthy donor and P937R mutated patients were already present in our laboratory.

**Table 3.**
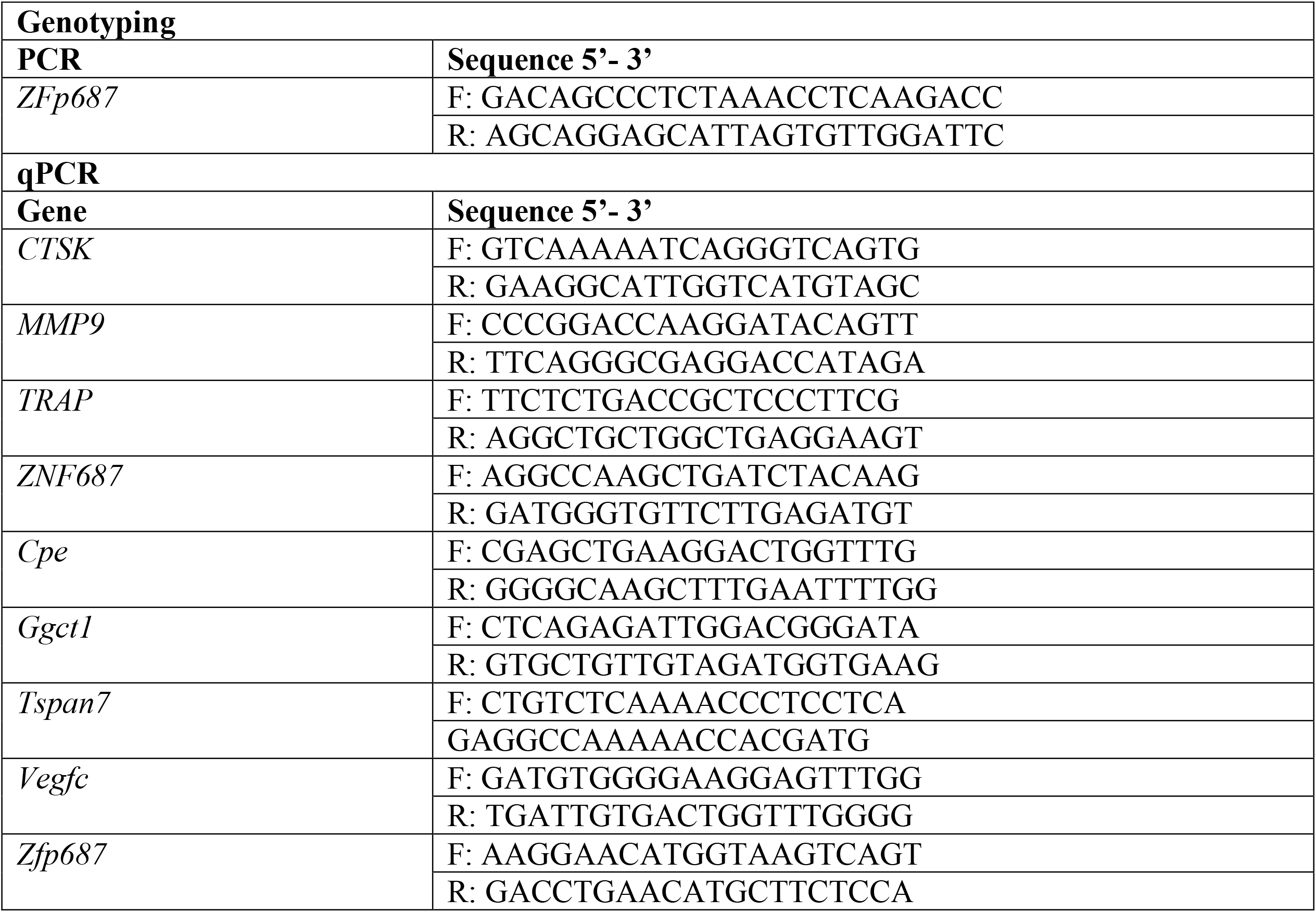
Sequence of primers used for genotyping and qRT-PCR:

### Library preparation and RNA-sequencing

The libraries were generated using depleted RNA obtained from 1 μg of total RNA by TruSeq Sample Preparation RNA Kit (Illumina, Inc., San Diego, CA, USA) according to the manufacturer’s protocol without further modifications. All libraries were sequenced on the Illumina HiSeq 1000, generating 100 bp paired end reads. Illumina BCL2FASTQ v2.20 software was used for de-multiplexing and production of FASTQ sequence files. FASTQ raw sequence files were subsequently quality checked with FASTQC software (http://www.bioinformatics.bbsrc.ac.uk/projects/fastqc). Subsequently, sequences with low quality score or including adaptor dimers or mitochondrial or ribosomal sequences, were discarded from the analysis. The resulting set of selected reads were aligned onto the complete Mouse genome using Spliced Transcripts Alignment to a Reference algorithm STAR version 2.7.3 using GRCm39 Genome Assembly and GRCm39.105.gtf as gene definition^43^. The resulting Mapped reads were used as input for feature Counts function of Rsubread packages and used as Genes counts for Differentially expression analysis using Deseq2 package^44^. We used shrinkage estimator from the apeglm package for visualization and ranking^45^. Differentially expressed genes (DEGs) were selected based on adjusted p-value < 0.05 and by setting Log2FoldChange ≥ 1 for upregulated and ≤ −1 for downregulated genes. The cut-off was further increase to Log2FoldChange ≥ 2 for upregulated and ≤ −2 for downregulated genes for a more stringent filter. To select genes that remained unchanged in wild type osteoclasts we selected DEGs genes that showed an Absolute Log2FoldChange ≤ 0.5 in *Zfp687*^+/-^ osteoclasts analysis. Selected DEGs were used as input to perform pathway enrichment analysis by IPA system (Ingenuity® Systems, www.ingenuity.com). IPA annotation was used as starting point for focusing in osteoclast differentiation.

### Quantification and statistical analyses

All data are presented as mean or median ± standard deviation (s.d.). The sample size for each experiment and the replicate number of experiments are included in the figure legends. Statistical significance was defined as p < 0.05. Statistical analyses were performed using ordinary one-way ANOVA or student’s-T-test (GraphPad Prism; version 9.3.1).

---

**Figure S1.**
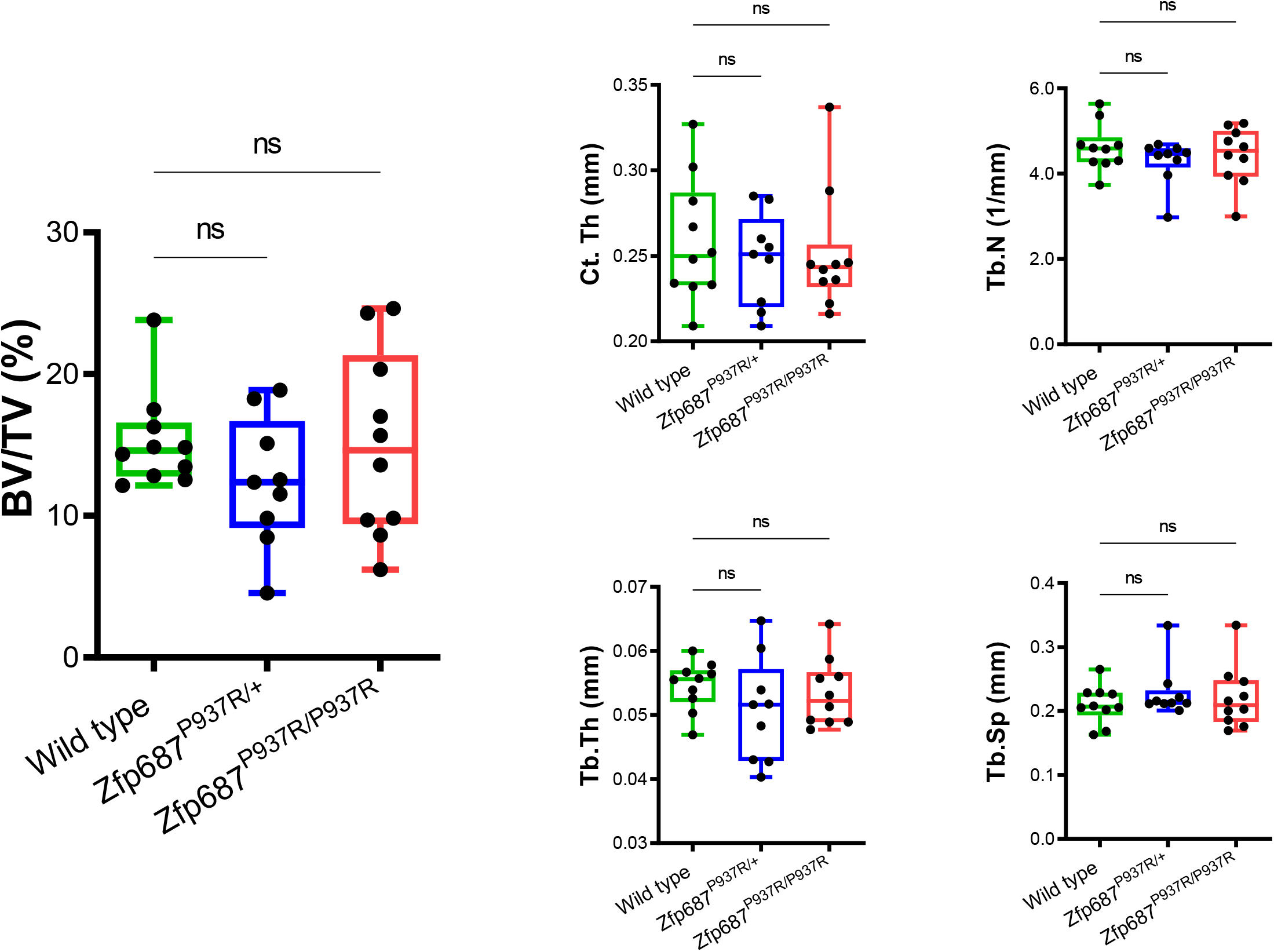

**Figure S2.**
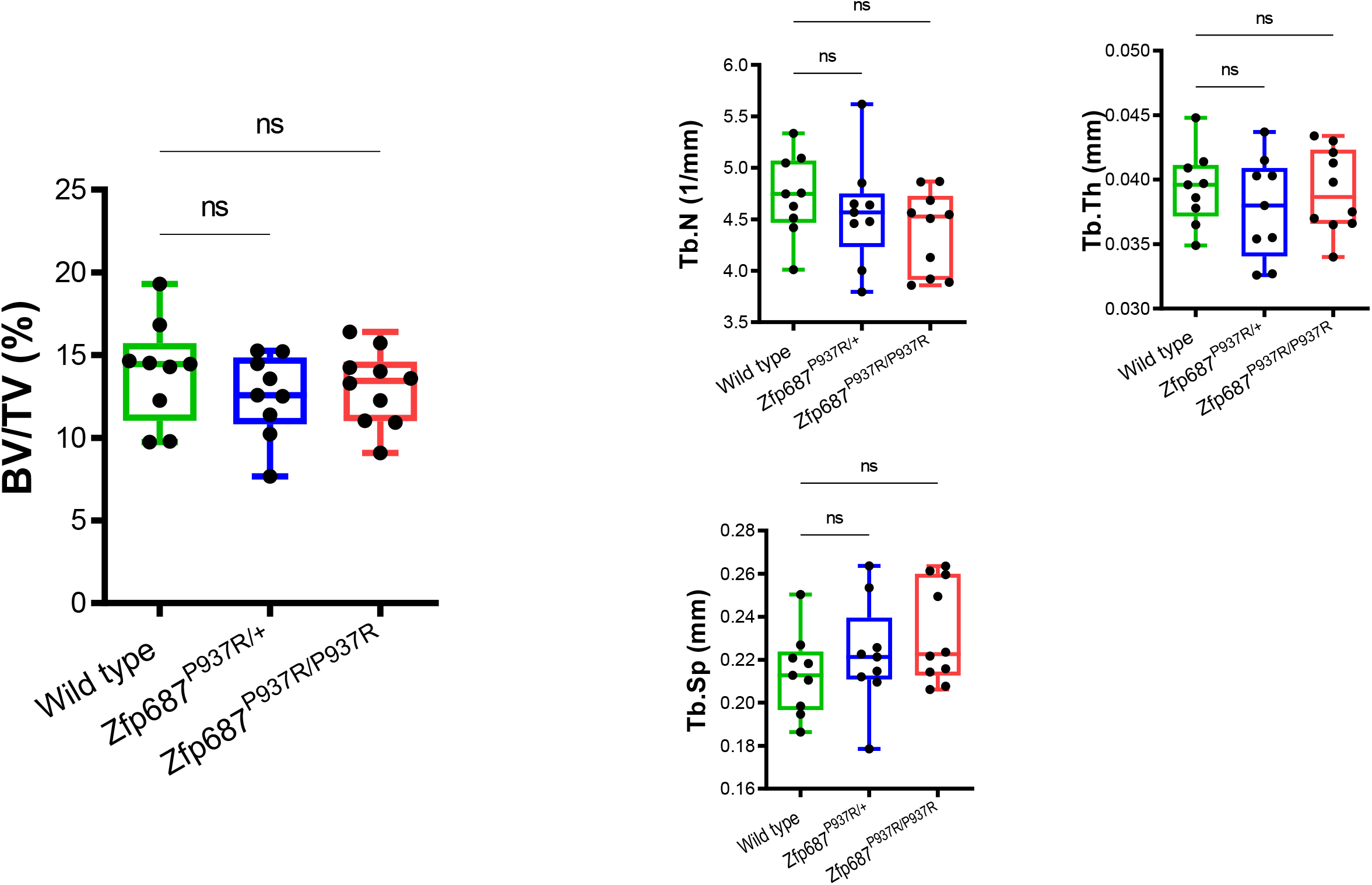

**Figure S3.**
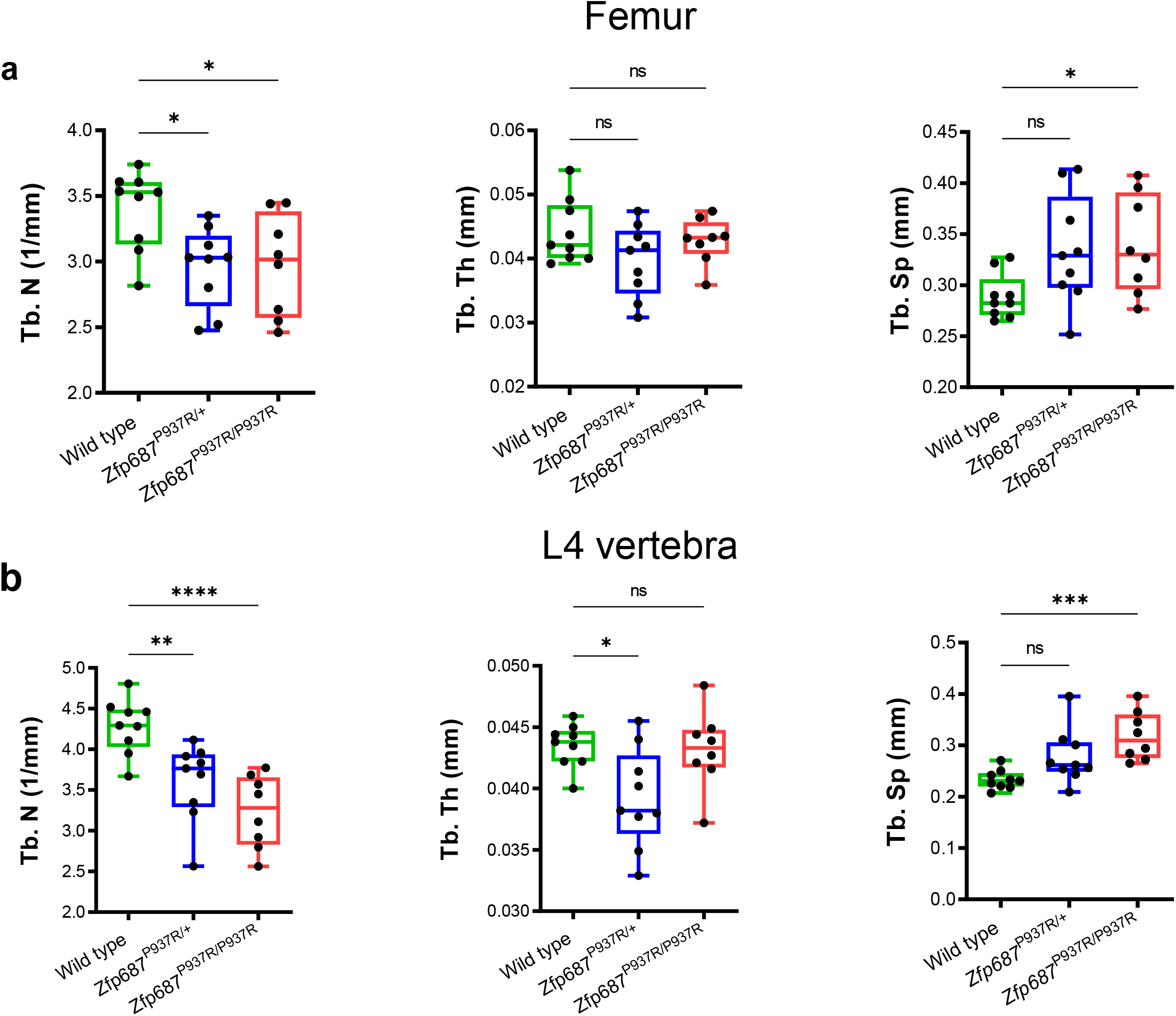

## Notes

### Competing Interest Statement

The authors have declared no competing interest.

